# A validated pangenome-scale metabolic model for the *Klebsiella pneumoniae* species complex

**DOI:** 10.1101/2023.12.20.572682

**Authors:** Helena B. Cooper, Ben Vezina, Jane Hawkey, Virginie Passet, Sebastián López-Fernández, Jonathan M. Monk, Sylvain Brisse, Kathryn E. Holt, Kelly L. Wyres

## Abstract

The *Klebsiella pneumoniae* Species Complex (KpSC) is a major source of nosocomial infections globally with high rates of resistance to antimicrobials. Consequently, there is growing interest in understanding virulence factors and their association with cellular metabolic processes for developing novel anti-KpSC therapeutics. Phenotypic assays have revealed metabolic diversity within the KpSC, but metabolism research has been neglected due to experiments being difficult and cost-intensive.

Genome-scale metabolic models (GSMMs) represent a rapid and scalable *in silico* approach for exploring metabolic diversity, which compiles genomic and biochemical data to reconstruct the metabolic network of an organism. Here we use a diverse collection of 507 KpSC isolates, including representatives of globally distributed clinically-relevant lineages, to construct the most comprehensive KpSC pan-metabolic model to-date, KpSC pan v2. Candidate metabolic reactions were identified using gene orthology to known metabolic genes, prior to manual curation via extensive literature and database searches. The final model comprised a total of 3,550 reactions, 2,403 genes and can simulate growth on 360 unique substrates. We used KpSC pan v2 as a reference to derive strain-specific GSMMs for all 507 KpSC isolates, and compared these to GSMMs generated using a prior KpSC pan-reference (KpSC pan v1) and two single-strain references. We show that KpSC pan v2 includes a greater proportion of accessory reactions (8.8%) than KpSC pan v1 (2.5%). GSMMs derived from KpSC pan v2 also result in more accuracy growth predictions than those derived from other references in both aerobic (median accuracy = 95.4%) and anaerobic (median accuracy = 78.8%). KpSC pan v2 also generates more accurate growth predictions, with high median accuracies of 95.4% (aerobic, n=37 isolates) and 78.8% (anaerobic, n=36 isolates) for 124 matched carbon substrates.

KpSC pan v2 is freely available at https://github.com/kelwyres/KpSC-pan-metabolic-model, representing a valuable resource for the scientific community, both as a source of curated metabolic information and as a reference to derive accurate strain-specific GSMMs. The latter can be used to investigate the relationship between KpSC metabolism and traits of interest, such as reservoirs, epidemiology, drug resistance or virulence, and ultimately to inform novel KpSC control strategies.

**Significance as a BioResource to the community:** *Klebsiella pneumoniae* and its close relatives in the *K. pneumoniae* Species Complex (KpSC) are priority antimicrobial resistant pathogens that exhibit extensive genomic diversity. There is growing interest in understanding KpSC metabolism, and genome scale metabolic models (GSMMs) provide a rapid, scalable option for exploration of whole cell metabolism plus phenotype prediction. Here we present a KpSC pan-metabolic model representing the cellular metabolism of 507 diverse KpSC isolates. Our model is the largest and most comprehensive of its kind, comprising >2,400 genes associated with >3,500 metabolic reactions, plus manually curated evidence annotations. These data alone represent a key knowledge resource for the *Klebsiella* research community; however, our model’s greatest impact lies in its potential for use as a reference from which highly accurate strain-specific GSMMs can be derived to inform in depth strain-specific and/or large-scale comparative analyses.

**Data summary:**

1. *Klebsiella pneumoniae* species complex (KpSC) pan v2 metabolic model available at https://github.com/kelwyres/KpSC-pan-metabolic-model.
2. All KpSC isolate whole genome sequences used in this work were reported previously and are available under Bioprojects PRJEB6891, PRJNA351909, PRJNA493667, PRJNA768294, PRJNA253462, PRJNA292902 and PRJNA391323. Individual accessions listed in Table S1.
3. Strain-specific GSMMs used for comparative analyses (deposited in Figshare - 10.6084/m9.figshare.24871914), plus their associated MEMOTE reports (indicates completeness and annotation quality), reaction and gene presence-absence matrices across all isolates.
4. Growth phenotype predictions derived from strain-specific GSMMs are available in Table S4.
5. Binarised Biolog growth phenotype data for n=37 isolates (plates PM1 and PM2, aerobic and anaerobic conditions) are available in Tables S6 & S7.
6. Additional growth assay data for six substrates not included on Biolog plates PM1 and PM2 (deposited in Figshare - 10.6084/m9.figshare.24871914).

## 2. Introduction

*Klebsiella pneumoniae* and closely related organisms in the *K. pneumoniae* Species Complex (KpSC) are a major source of nosocomial infections globally and among the top three causes of deaths associated with antibiotic resistance (1). Consequently, the World Health Organization has designated development of novel anti-*Klebsiella* control strategies an urgent global priority (2). Many existing antimicrobials target or alter cellular metabolic processes, and recent works have highlighted a role for metabolism in *K. pneumoniae* virulence (3,4), supporting these processes as prime targets for novel therapeutics. Despite this, our understanding of KpSC metabolism remains comparatively limited. The matter is further complicated by metabolic variability (5,6), likely driven by substantial genetic variation: The *K. pneumoniae* population consists of hundreds of distinct ancestral lineages (‘clones’) where gene content varies significantly within and between clones (7). The total pangenome is estimated to exceed 100,000 protein coding sequences (8), with ∼37% of these predicted to be involved in metabolism. However, it can be difficult to interpret gene level information in the context of metabolic phenotypes, and large-scale laboratory growth phenotype experiments are infeasible due to their time and cost intensive nature, so the true extent of population variation remains unknown.

Genome-scale metabolic models (GSMMs) are a powerful *in silico* approach to explore the metabolism of individual cells, and predict metabolic phenotypes such as the ability to grow on a given substrate or the outcomes of gene knock-out mutations (Figure 1). Such analyses have been used to identify novel anti-*K. pneumoniae* drug targets (3,9) and indicate substrate preferences among clinical isolates (3,10). GSMMs are a mathematical representation of an individual strain’s metabolism derived from published literature, genome annotations and biochemical data (11,12). They contain stoichiometrically balanced reactions that are centred around the Biomass Objective Function, representing the growth requirements for producing a single cell (Figure 1). Gene information for every reaction is summarised as a Gene-Protein-Reaction (GPR) rule (Figure 1), which summarises the genes required to catalyse a reaction, along with their associated nucleotide and amino acid sequences. Reactions are assigned a confidence score based on the level of evidence available to support their occurrence in the species of interest, with protein characterisation being the strongest form of evidence (11).

**Figure 1:**
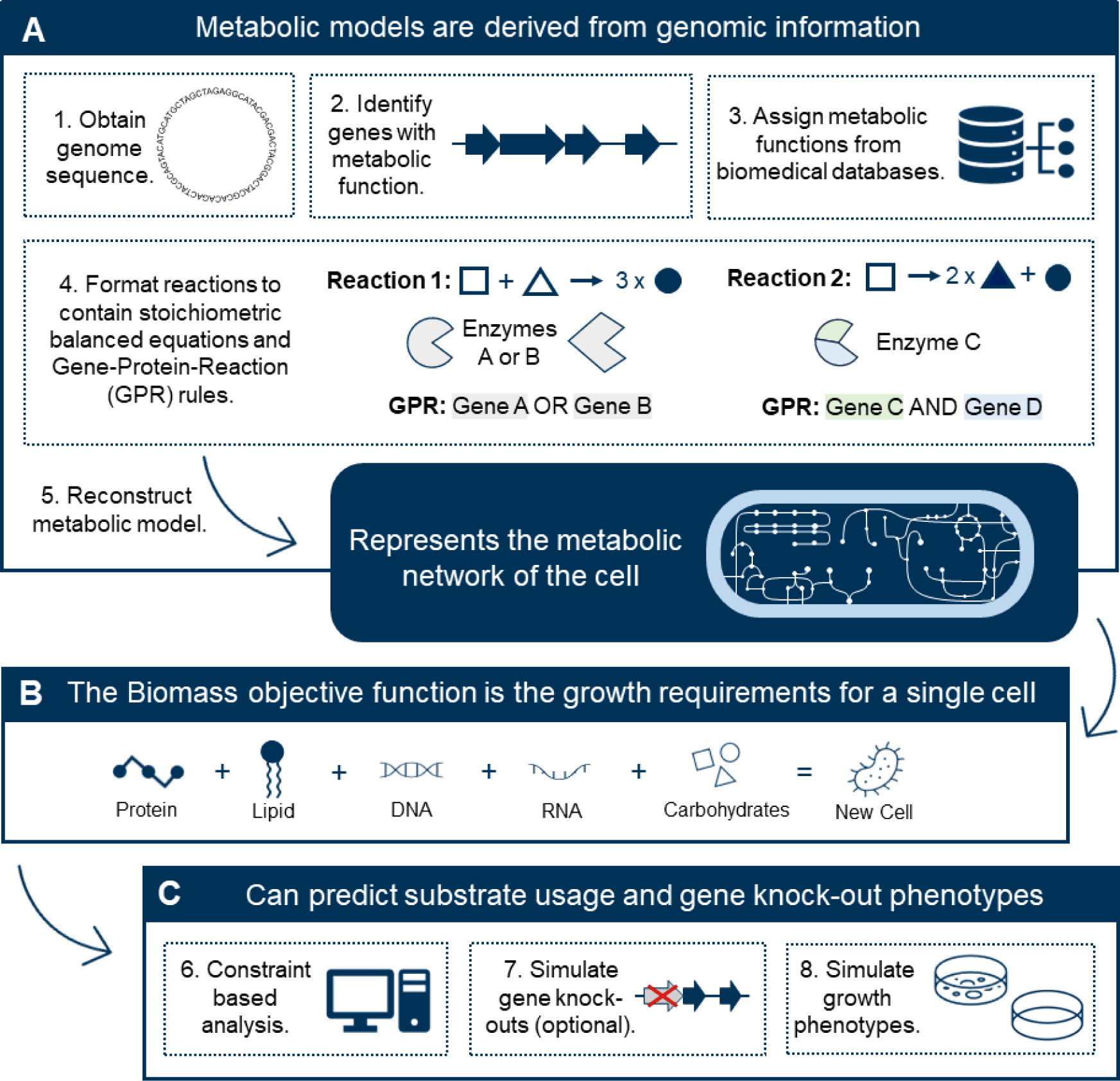
G**e**nome **scale metabolic model (GSMM) overview.** A) Summary of the typical approach for generating a GSMM. B) Constituent components of the Biomass Objective Function, which defines the requirements for production of new cells. Optimisation of the Biomass Objective Function via constraint based analysis (C), can be used to predict growth in different conditions i.e. where an optimised objective value ≥ 0.001 indicates growth, and an objective value < 0.001 indicates no growth. C) Common applications of GSMMs.

GSMMs can be built *de novo* by identifying candidate metabolic reactions from genome annotations and experimental data, before assembling and manually refining them into a draft model; this is a very time consuming process that is not feasible for large genome collections (11). An alternative approach leverages reference models as templates to extract strain-specific models (13). We recently implemented this approach in an automated pipeline Bactabolize, which enables rapid and scalable model generation and phenotype prediction (14). We also described a KpSC pan-reference model (KpSC pan v1) based on 37 strain-specific models (14), each derived from *K. pneumoniae* MGH 78578 model iYL1228 (15) with minimal manual curation (6). Using Bactabolize and KpSC pan v1, we showed that the reference based approach can generate highly accurate strain-specific models that equalled or surpassed existing scalable model generation methods (14). However, models generated for KpSC clones not represented in the KpSC pan v1 model tended to have a lower accuracy compared to those that were present (14), indicating room for improvement.

In this study, we generated a novel KpSC pan-reference model using a large, diverse collection of 507 KpSC isolates, which captures more metabolic information than the prior pan model. When used as a reference, KpSC pan v2 resulted in a more diverse collection of strain-specific GSMMs, with higher growth prediction accuracies as validated with phenotype data. Our data highlights the value of pan-metabolic reference models to support large-scale comparative metabolic modelling analyses of diverse bacterial species, such as *K. pneumoniae* and other priority antimicrobial resistant pathogens.

## 3. Methods

### Genome collection and pangenome construction

Figure 2 summarises our approach for developing the KpSC pan v2 reference model. A starting dataset of 510 *Klebsiella* genome assemblies were collected (Table S1), comprising 452 isolate genomes from the *Klebsiella* Acquisition Surveillance Project at Alfred Health (KASPAH) (16), a diverse collection for which all isolates were available in house for phenotypic validation. In addition, 58 genomes for which draft metabolic models were published previously (n=20/22 multi-drug resistant *K. pneumoniae* (17), n=37 diverse KpSC (6) and n=1 hypervirulent *K. pneumoniae* (18)), were also included. KASPAH isolates were *de novo* assembled using SPAdes optimised with Unicycler v0.4.7 as reported previously (16,19) and all remaining assemblies were retrieved from public databases (5,6,17,18). To ensure the assemblies were of sufficient quality to produce accurate metabolic models, we applied a tiered quality control framework as described previously (14). Genomes were filtered using a threshold of ≤200 assembly graph dead-ends (where assembly graphs were available, n=452 KASPAH genomes), and subsequently by N50 ≥65,000 bp (Table S1). The 507 genomes that passed quality control were annotated using prokka v1.13.3 (20) with the following parameters: --gcode 11, --addgenes and --protein to initially annotate proteins using *K. pneumoniae* HS12286 (21). These were then compiled into a pangenome using panaroo v1.1.2 (22) (default parameters, except for protein family sequence identity and length difference, both of which were set to 90%). Kleborate v2.3.2 was used to determine the seven gene multi-locus sequence types (STs), antibiotic resistance determinants and capsule types (23,24).

**Figure 2:**
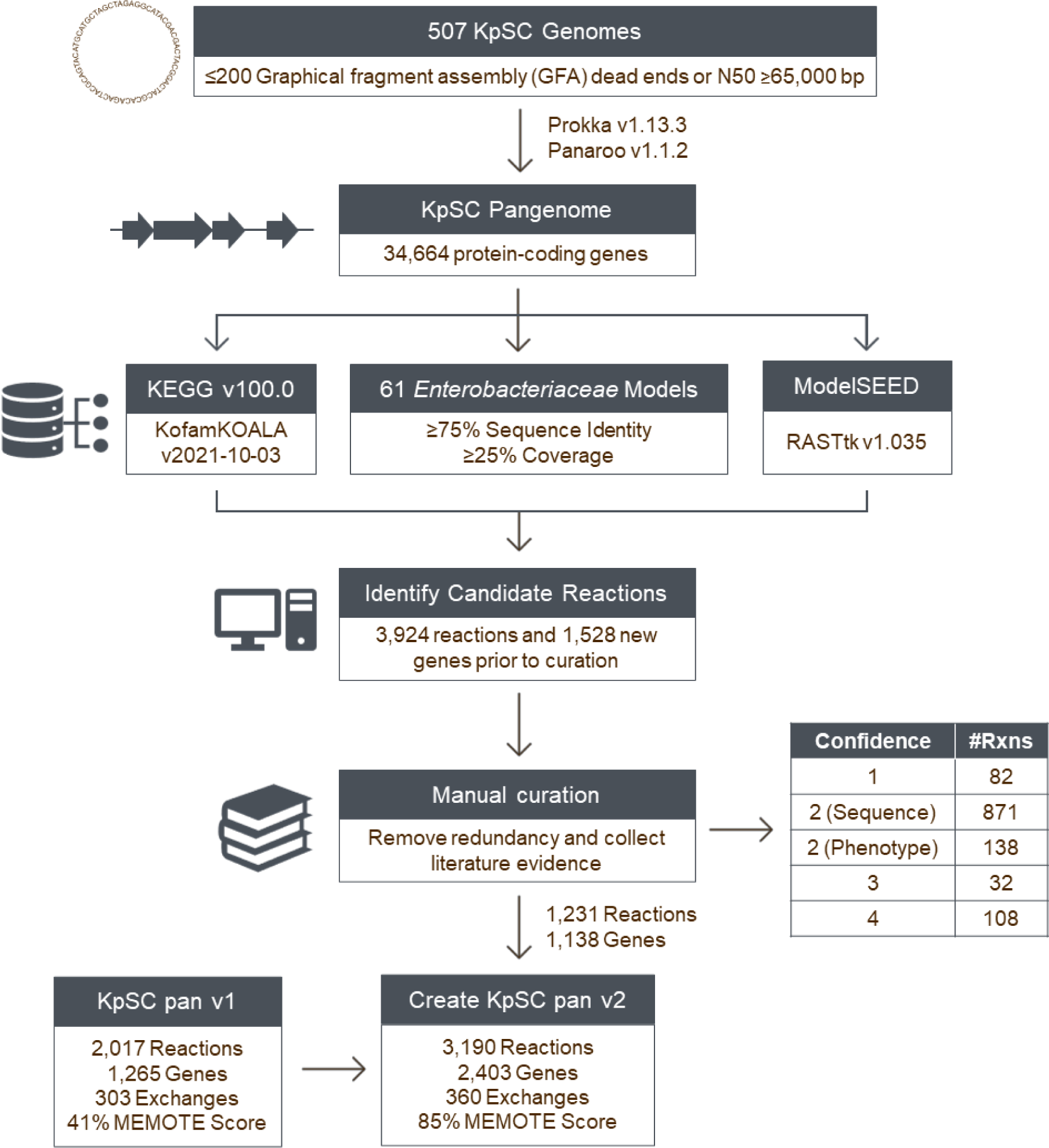
Approach for generating KpSC pan v2. Blue boxes indicate major analysis steps and data sources, whereas white boxes indicate additional information. The table adjacent to manual curation is a breakdown of the evidence confidence scores (Confidence) and the number of reactions (#Rxns) in each category (12). The final set of curated gene and reaction information is available in Table S3 and summarised in Table 1.

### Identification and curation of metabolic reactions

Representative sequences from each of the pangenome ortholog clusters were used to find new candidate metabolic reactions based on sequence homology to proteins identified from three sources (Figure 2):

1. **Published *Enterobacteriaceae* GSMMs:** We identified 61 published metabolic models for which corresponding sequence data were publicly available (Table S2), including 50 *Escherichia coli* (25–28), eight *Shigella* (28), one *Yersinia pestis* (29) two *Salmonella enterica* (30,31) and five additional *K. pneumoniae* models (6,17,18). Models were downloaded from BiGG (32), BioModels (33) and K-Base (34); genomes were downloaded from NCBI (35). Sequence orthologs were identified by BLASTn and BLASTp bidirectional best hit (BBH) search with coverage ≥25% and identity ≥75% as described previously (6,13,36). Reactions associated with the BBH were subject to manual curation as described below. Reactions for which the GPR comprised n>1 required genes (i.e. Gene X AND Gene Y) were considered for curation if n-1 gene orthologs were identified within the KpSC pangenome with identity ≥75%. Additionally, reactions matching those in the existing KpSC pan v1 model for which novel orthologs were identified at <75% identity were considered for curation via update of the GPR to include the novel divergent ortholog as additional gene sequence variants.
2. **KEGG:** Metabolic reactions were identified from KEGG v100.0 (37) using kofamscan v2021-09-05 (38) (significance value of ≤0.001).
3. **ModelSEED:** RASTtk 1.3.0 was used to assign enzyme commission numbers to each representative KpSC pangenome sequence, and these were matched back to the ModelSEED v2.6.1 database to extract the corresponding metabolism and reaction data (39,40).

Candidate reactions were filtered to remove redundancy by; i) comparing their associated protein sequences (80% identity threshold); ii) merging reactions identified from multiple sources; and iii) removing information already captured by the KpSC pan v1 model. The remaining reactions were manually curated to ensure they had sufficient evidence for inclusion in the KpSC pan v2 model via a literature search (11) and were formatted according to COBRApy conventions (41). Each reaction was given a confidence score based on the evidence supporting its inclusion in the model; e.g. a score of two corresponds to sequence homology evidence alone and/or phenotypic evidence that *Klebsiella* can metabolise the associated substrate(s) (11). Documented gene knock-out data showing that the absence of a gene causes a loss in phenotype is considered higher level evidence (score 3), as is a documented solved *Klebsiella* protein structure (score 4) (11).

The final set of curated reactions (Table S3) and associated sequence data were combined with those from the KpSC pan v1 model to create the updated KpSC pan-reference model, known as KpSC pan v2.

### Strain-specific metabolic models and growth phenotype predictions

In order to assess the metabolic content and diversity represented by the KpSC pan v2 model, we compared its performance as a reference model against those of the KpSC pan v1 model (14) and two curated single-strain *K. pneumoniae* models (iYL1228 (15) and iKp1289 (18)). We used Bactabolize v1.0.2 (14) to build four independent strain-specific GSMMs and independently predict growth phenotypes for each of the 507 genome assemblies that passed quality control. All possible sources of carbon, nitrogen, sulfur and phosphorus substrates were tested as the sole source of the respective element in M9 minimal media, in both aerobic and anaerobic conditions. Prior to use as a reference, iYL1228 was updated to add the chemical formulas for every metabolite using BiGG (32), which was required to allow Bactabolize to detect the potential growth sources (14). A sink reaction in iKp1289 (sink DASH dna5mtc_c0) was also modified to be non-reversible to ensure methylated DNA was not being used as a carbon source (18). Draft models were built using the draft_model command (default parameters) and growth phenotype predictions were performed using the fba command (default parameters except; fba_open_value = -20). When iKp1289 was used as a reference, media_type = m9_SEED_media and fba_spec_name = m9_SEED_spec were set as iKp1289 uses ModelSEED IDs rather than BiGG IDs. Simulated growth phenotypes with a predicted biomass value >0.001 were considered positive for growth and those with biomass ≤0.001 were considered negative (Table S4). Comparisons to other metabolic modelling pipelines, such as CarveMe (42), were not performed in this study as the KpSC pan v1 model has been previously benchmarked against several pipelines and was shown to outperform or equal these (14).

### Estimating model accuracies

To determine the accuracy of the KpSC pan v2 model, we collected two types of *in vitro* data, the first being Biolog growth phenotypes for 190 carbon substrates (PM1 and PM2 plates) in aerobic and anaerobic conditions (for n = 37 KpSC isolates). The second set of data were derived from independent growth assays for six additional carbon substrates tested in aerobic conditions only (n = 37 to 42 KpSC isolates per substrate, Table S5).

The aerobic Biolog data has been described previously by Blin *et al.* (5), with the anaerobic data generated via the same protocol with the following differences: Strains were cultured onto Luria-Bertani agar at 37°C for 48 hours within an anaerobic chamber. All required reagents and materials were placed inside an anaerobic chamber for 48 hours in parallel. Biolog 96-well plates were placed in anaerobic jars containing 2.5 L anaeropack catalyzers (Thermo scientific, France) for 48 hours before being transferred to the anaerobic chamber, where the remainder of the experiment took place. For each strain, IF-0a fluid was inoculated with fresh colonies to reach 40% transmittance. The bacterial preparation was diluted at 1/20 in IF-0a, and then to 1/5 in a solution containing tetrazolium chloride (Dye D), menadione, potassium ferricyanide and water.

The aerobic growth threshold was defined empirically (5), with a maximum value greater than 150 on the respiration curve indicating positive growth. To determine the anaerobic growth threshold, we plotted the distribution of the maximum values from the Biolog respiration curve for every substrate and isolate tested to identify the inflection point between the bimodally distributed data (i.e. growth cut off points). The distribution was distinct from that of the aerobic data (5), and we hypothesised that this may be driven by species-specific variations (Figure S1). Plotting the distributions by species confirmed our hypothesis (Figure S1), and informed species-specific thresholds for *K. pneumoniae* (maximum value > 165), *Klebsiella quasipneumoniae* subsp. *quasipneumoniae* (maximum value > 155), *Klebsiella quasipneumoniae* subsp. *similipneumoniae* (maximum value > 150) and *Klebsiella variicola* subsp. *variicola* (maximum value > 160). As *Klebsiella africana, Klebsiella variicola* subsp. *tropica* and *Klebsiella quasivariicola* were only represented by one isolate each, we could not determine species-specific thresholds and therefore used a threshold of 190, derived from the combined distribution (Figure S1).

Independent phenotype assays were performed for six substrates not present on the Biolog PM1 and PM2 carbon plates, which were chosen because they were newly implemented in the KpSC pan v2 model (either via addition of new exchange reactions or by completion of a metabolic pathway that was partially present in KpSC pan v1), they were available for purchase and import to Australia, and were not prohibitively expensive. All substrates were obtained from Sigma-Aldrich: uracil (CAS 66-22-8), allantoin (CAS 97-59-6), 2- aminoethylphosphonic acid (CAS 2041-14-7), L-ectoine (CAS 96702-03-03), hydrocinnamic acid (CAS 501-52-0) and beta-alanine (CAS 107-95-9). Growth assays were performed for the 37 KpSC isolates with existing Biolog data (5) and up to five additional isolates (16,43) to confirm the predicted growth variability across the wider population. The protocol was described previously by Hawkey *et al.* (6). Briefly, isolates were grown overnight in 5 mL of minimal media (2x M9, Minimal Salts [Sigma-Aldrich], 2 mM MgSO4 and 0.1 mM CaCl2) and 20 mM D-Glucose (CAS 50-99-7) at 37°C, shaking at 200 rpm. Overnight cultures were diluted to the McFarland standard of 0.4-0.55 turbidity. Dilutions of 20 μL were inoculated in triplicate to 96-well cell culture plates (Corning) containing M9 minimal media with either 20 mM substrate (4mM for allantoin due to low solubility (44)), Milli-Q H2O (negative control) or 20 mM D-glucose (positive control) at pH 7.0 (pH 6.0 for uracil to avoid precipitation). Plates were incubated at 37°C and the OD600 values recorded after 24 and 48 hours using the Infinite 200 PRO plate reader (Tecan) using Tecan i-control version 2.0.10.0, firmware V_4.31_06/19_Infinite, at 600 nm absorbance after 30 seconds of shaking at 218.3 rpm (amplitude of 3mm). Isolates were marked as positive for growth if the adjusted OD600 fold change (calculated as indicated below) exceeded the substrate-specific threshold (determined from the empirical distributions, see Figure S2). Positive growth and fold-change were determined using the following formulae where TR refers to Technical Replicates:

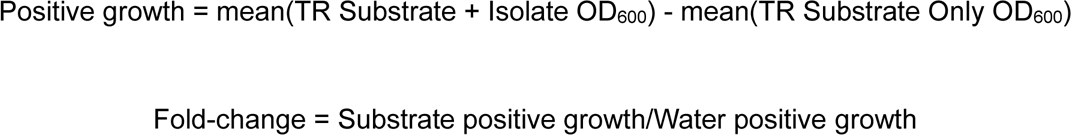

We additionally tested two predicted L-histidine auxotrophs and two control isolates for growth on L-histidine (Sigma-Aldrich, CAS 71-00-1). Experiments were performed as described above with the exception that the primary overnight cultures were grown in M9 minimal media plus 20mM D-glucose and 20mM L-histidine. Residual histidine was removed by centrifugally washing overnight cultures twice with 1.5 mL 0.9 M NaCl at 8000 x g for 15 minutes, then a final resuspension in 0.9 M NaCl prior to dilution to the McFarland standard of 0.4-0.55.

*In vitro* data were compared to *in silico* predictions; accuracy, sensitivity and specificity were calculated for the strain-specific GSMMs derived from each reference model using the following formulae where TP = True Positive, TN = True Negative, FP = False Positive, FN = False Negative:

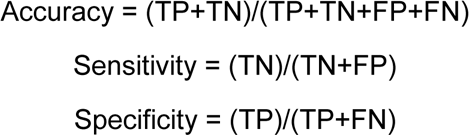

These metrics were calculated using multiple subsets of substrates so that direct comparisons could be made between models generated using different references, which each support different substrates (see Tables S6 & S7). The first comparison was for the set of 89 substrates that could be simulated by all models. Additional comparisons represented the set of 91 substrates that could be simulated by all models except those generated using the iKp1289 reference, and the set of 105 substrates that could be simulated by all models except those generated using iKp1289 and iYL1228. Final accuracy for KpSC pan v2 was calculated for the maximum number of substrates with matched *in vitro* and *in silico* data (n=124).

## 4. Results

### Diverse collection of isolate genomes

The initial dataset of 510 isolates contained at least one representative from each of seven of the KpSC subspecies (including 79.0% *K. pneumoniae*, 15.1% *K. variicola* & 5.3% *K. quasipneumoniae,* Table S1), 268 unique STs and 115 distinct capsule polysaccharide loci (Table S1), notably including representatives of 12 of the 13 global problematic clones (7). The dataset was diverse in terms of isolate sources, including 344 hospital-acquired infection, 45 community-acquired infection, 126 human gut carriage and 13 environmental isolates, with 23.7% being multidrug resistant (acquired resistance to at least three antimicrobial classes, including 13.7% extended-spectrum beta-lactamases, 6.1% carbapenemases) (Table S1).

### Description of KpSC pan v2

Rather than building the reference model *de novo*, we used the KpSC pan v1 model (14) as the starting point to incorporate additional metabolic reactions and/or divergent gene orthologs. Candidate orthologs and their associated metabolic reactions were identified by comparing representative gene sequences derived from the pangenome of 507 KpSC isolates (excludes three genomes which did not pass quality control, see Methods) to those of published *Enterobacteriaceae* GSMMs (18,25–31), KEGG (37) and ModelSEED (39) databases (Figure 2). This pangenome contained 34,664 unique genes with 11.4% core (present ≥95% of isolates) and 88.6% accessory (present in <95% of isolates). Following manual curation (summarised in Table S3), 1,141 gene sequences were confirmed as orthologs, corresponding to 1,058 unique metabolic reactions (883 catalytic, 175 transport) that were not present in KpSC pan v1. These reactions were formatted as new GPR rules and added to the KpSC pan v2 model along with 173 reactions (18 catalytic, 63 transport, 92 pseudo) that were not associated with a GPR rule.

Most novel GPR-associated reactions were identified from *Enterobacteriaceae* GSMMs only (44.9%) or KEGG only (46.8%). The remaining reactions were identified from *Enterobacteriaceae* GSMMs combined with information from KEGG (6.9%) or ModelSEED (0.2%), and ModelSEED only (1.2%). Each reaction was curated and assigned a confidence score (11), majority with confidence score two (81.9%) which corresponded to 70.7% genomic and 11.2% phenotypic evidence (Table S3). The remaining reactions comprised 8.8% confidence score four (e.g. protein structures), 2.6% confidence score three (e.g. gene knock-outs) and 6.7% confidence score one (i.e. required for modelling but without additional supporting evidence). Of these curated reactions, 50.1% were supported by evidence from *Klebsiella,* with the remaining supported by evidence from other *Enterobacteriaceae* or other bacterial species (all confidence score ≤2, Table S3).

As an example of curation, a candidate reaction that transports periplasmic lipid A to the extracellular environment (14DENLIPIDAt2ex) requires the products of two genes for each subunit of the LptDE lipopolysaccharide transporter (45). We successfully identified an ortholog of one of the genes among our KpSC pangenome (>75% identity threshold), but the best match for the second gene (KpSC KPN_00673) had a sequence identity of just 70.4% when compared to that in *E. coli*. Following a literature search, we identified a published *K. pneumoniae* LptDE protein structure (45), and subsequently confirmed that the corresponding amino acid sequence shared 97.2% identity to KPN_00673, supporting the inclusion of the latter in the GPR rule.

The 92 novel pseudo-reactions without a GPR rule corresponded to: 85 exchange reactions, which enable the model to import/export metabolites into the cell; plus seven sink reactions which were required for the model to simulate growth on certain substrates for which the true degradation/export pathways have not been defined (Table S3). There were also 392 reactions (377 catalytic and 15 transport) that already existed in the KpSC pan v1 model that were updated to include additional gene sequence variants in their GPR rule. These changes included 21 reactions that were identified to contain erroneous GPRs inherited from previous *Klebsiella* models, such as a formamidase reaction (FORAMD) which was initially associated with a gene predicted to encode an oxamate carbamoyltransferase (Table S3).

To confirm that the KpSC pan v2 model was biologically viable, we used it as a reference to construct draft strain-specific models for our 507 KpSC isolates and simulate growth on glucose in M9 minimal media conditions - an expected growth phenotype for all *Klebsiella.* During this process we discovered that some of the models were unable to simulate production of D-mannose 1-phosphate (man1p_c), which was inherited as a biomass requirement from the first *K. pneumoniae* GSMM, iYL1228 (15). However, D-mannose 1- phosphate is a variable capsule component in *Klebsiella* that is not produced by all strains (46), and hence should not be considered as a generalised component for biomass production. We therefore created a new biomass function without the requirement for D-mannose 1-phosphate (BIOMASS_Core_Feb2022). Following this update, all but four isolates were predicted to simulate growth on glucose. The exceptions included *K. variicola* KSB2_2A and *K. pneumoniae* INF263 for which the draft genome assemblies lacked the *hisA* (KPN_02480) and *plsC* (KPN_03436) genes respectively (Table S1), likely absent due to having incomplete assemblies. Two complete assemblies for *K. pneumoniae* INF018 and *K. variicola* INF232 appeared to lack the entire *his* operon required for the production of histidine (confirmed as a genuine absence by aligning the reads from these isolate genomes to the *his* operon from *K. pneumoniae* INF149). *In vitro* experiments confirmed that INF018 and INF232 were histidine auxotrophs (Figure S3) and these isolates were excluded from further analyses.

The final KpSC pan v2 model contains 3,550 reactions (including 360 exchange reactions that define the set of substrates for which growth can be simulated) and 2,403 genes, an increase of 46.5% to 61.7% reactions and 86.4% to 95.5% genes, compared to the iYL1228, iKp1289 and KpSC pan v1 models (Table 1). For exchange reactions however, there was only an increase of 24.6% compared to the original iYL1228 model (Table 1), reflecting the limited *Klebsiella* growth phenotype data that is available to inform the addition of these exchanges. Nonetheless, the KpSC pan v2 model supports the prediction of 345 carbon, 192 nitrogen, 68 phosphorus and 34 sulphur growth sources in aerobic and anaerobic conditions (Table 1). However, 44 supported growth phenotypes were not considered biologically plausible, such as the use of the carbon component of secreted carbon dioxide as a sole carbon source (Table S4). The KpSC pan v2 model predicts the greatest number of growth phenotypes for sulfur and phosphorus utilisation, whereas it is the second highest to iKp1289 for carbon and nitrogen utilisation (Figure 3). Finally, the KpSC pan v2 model had the highest MEMOTE score of the four models tested (85%, Table 1), which indicates that the model is well annotated and stoichiometrically balanced (47).

**Figure 3:**
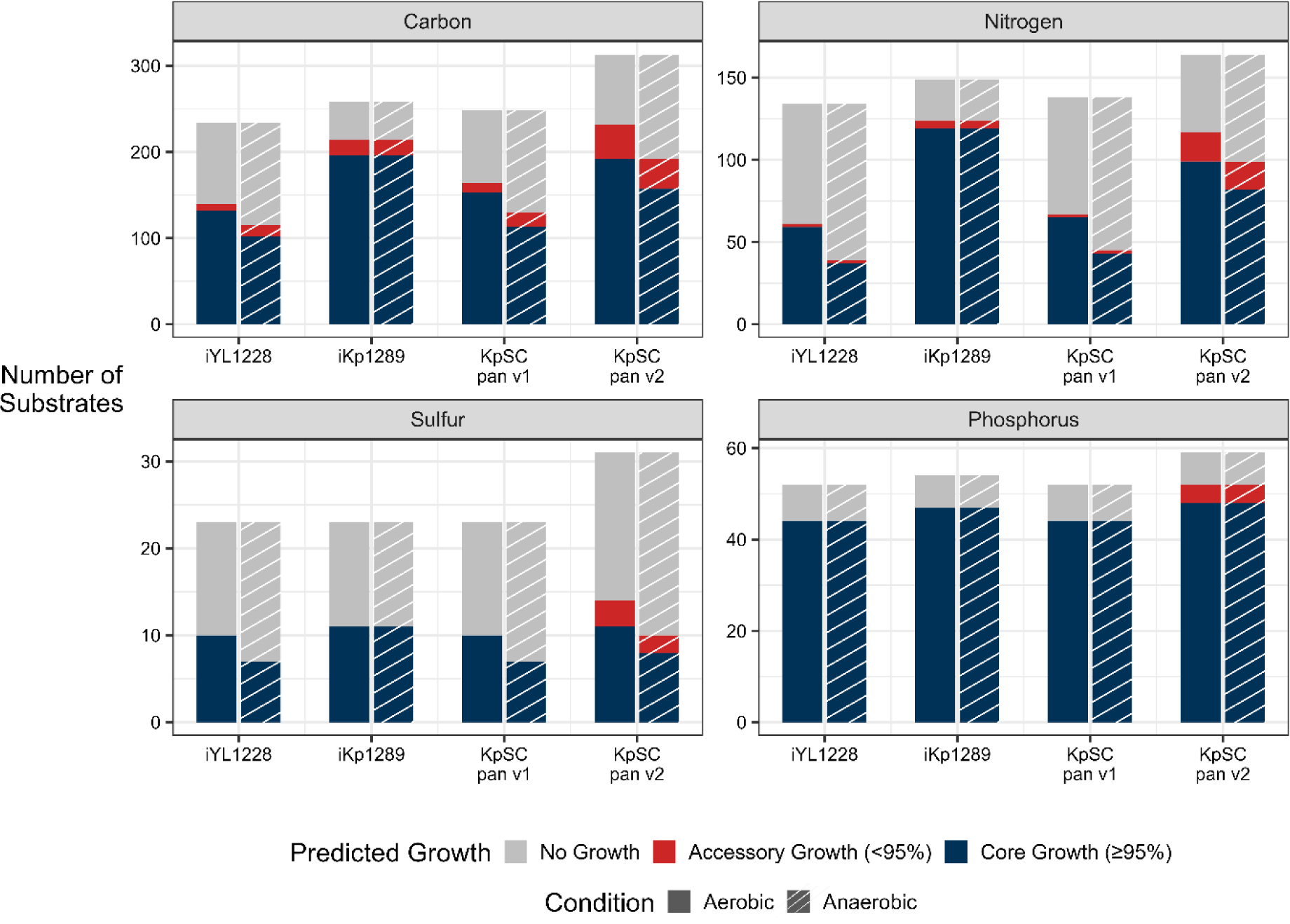
Growth simulation outcomes for strain-specific genome scale metabolic models derived from four different references. Simulations represent growth in minimal media supplemented with a single source of carbon, nitrogen, sulfur or phosphorus as indicated. The total number and set of substrates simulated is dependent upon those that can be supported by the reference model (see Table S4, note that substrates supported by the iYL1228 reference are a subset of those supported by KpSC pan v1, which are a subset of those supported by KpSC pan v2). Substrates supported by iKp1289 are an independent and overlapping set. Core growth (blue) refers to substrates for which growth was predicted for ≥95% of the population, whereas accessory growth (red) refers to substrates for which growth was predicted for <95% of the population. “No growth” (grey) refers to metabolites that are supported by the model, but do not result in positive growth predictions for any strains. These can occur due to incomplete metabolic pathways in the model (eg: the carbon utilisation pathway is complete but the nitrogen pathway is not, resulting in no growth on nitrogen) or where *Klebsiella* is legitimately unable to utilise the substrate for growth.

**Table 1:**
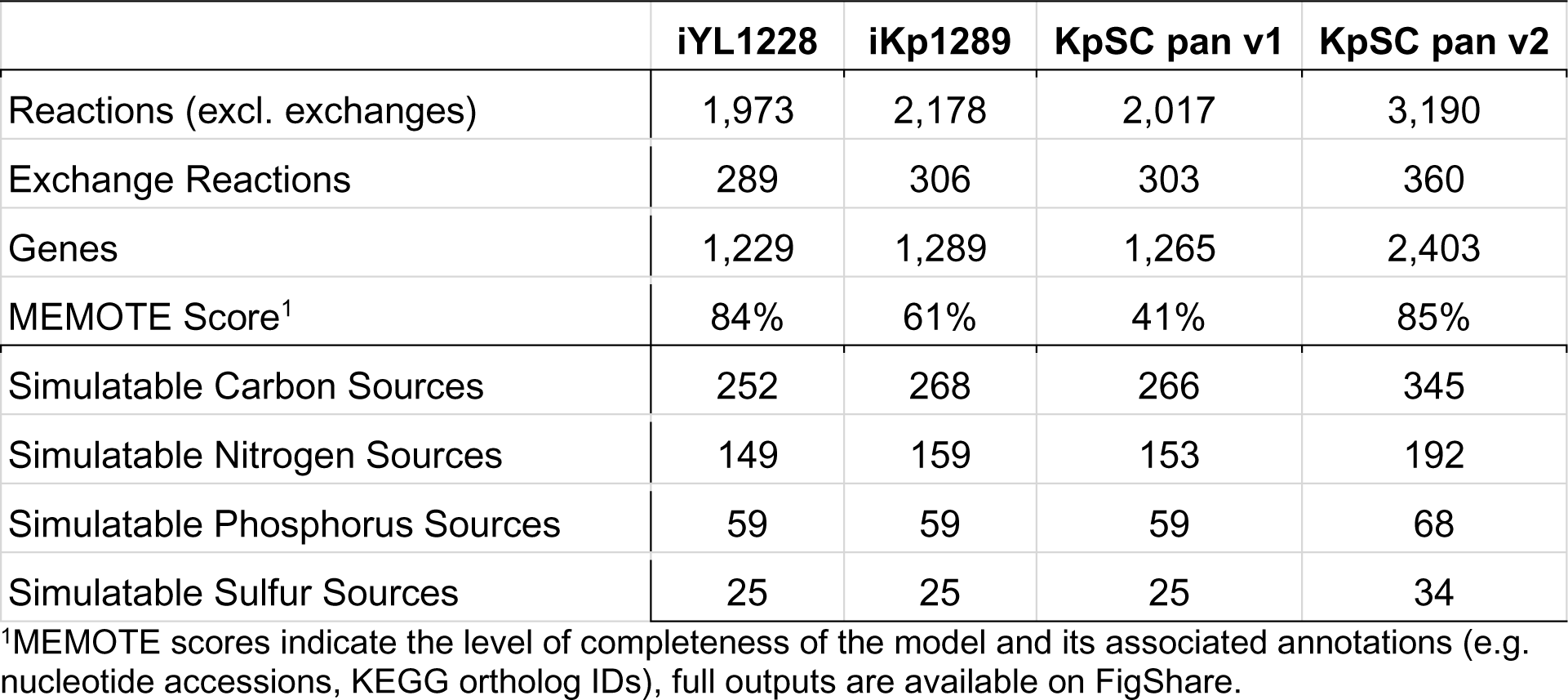
Features of KpSC pan v2 and previously published KpSC metabolic models.

### KpSC pan v2 captures the greatest metabolic variability

We compared strain-specific GSMMs and associated growth predictions generated using each of the four reference models (n = 505 KpSC isolates): iYL1228 (15), iKp1289 (18), KpSC pan v1 (14) and KpSC pan v2. The investigation of core (≥95% of isolates) and accessory (<95% of isolates) reactions was only performed for KpSC pan v1 (14) and v2, since iYL1228 was the direct predecessor of the KpSC pan v1 (6) and we were unable to satisfactorily harmonise the iKp1289 reaction nomenclature. Within the KpSC pan v1 model, 2,263 of 2,320 reactions (97.5%) were considered core, while 56 (2.5%) were accessory. In contrast, 3,236 of 3,550 reactions (91.2%) in the KpSC pan v2 model were core and 314 (8.8%) were accessory. Grouping reactions by KEGG subsystems (37) highlighted those associated with greater population variation with 28.8% (30/104) of xenobiotic biodegradation reactions in KpSC pan v2 considered accessory, as were 19.9% (87/437) carbohydrate metabolism reactions, indicating that KpSC pan v2 captures more accessory reactions than KpSC pan v1 (Figure S4).

Pairwise Jaccard gene distances (Figure 4A) showed that models built using the KpSC pan v2 reference capture greater strain-specific variation compared to those built using the KpSC pan v1 or single-strain references (median pairwise distances 0.053, versus 0.013 (iYL1228), 0.023 (iKp1289) and 0.020 (KpSC pan v1), respectively, p < 2.2×10^-16^ for all distribution comparisons by Mann-Whitney test). Two distinct peaks were also observed in the KpSC pan v2 Jaccard distributions (Figure 4), with the first peak corresponding to comparisons between isolates of the same species and the second to isolates of different species. This trend can also be seen to some extent in the KpSC pan v1 distribution but is notably absent from the single-strain references (Figure 4), highlighting that the pangenome approach has allowed the model to capture differences between species that would otherwise not be captured by a single-strain reference.\

**Figure 4:**
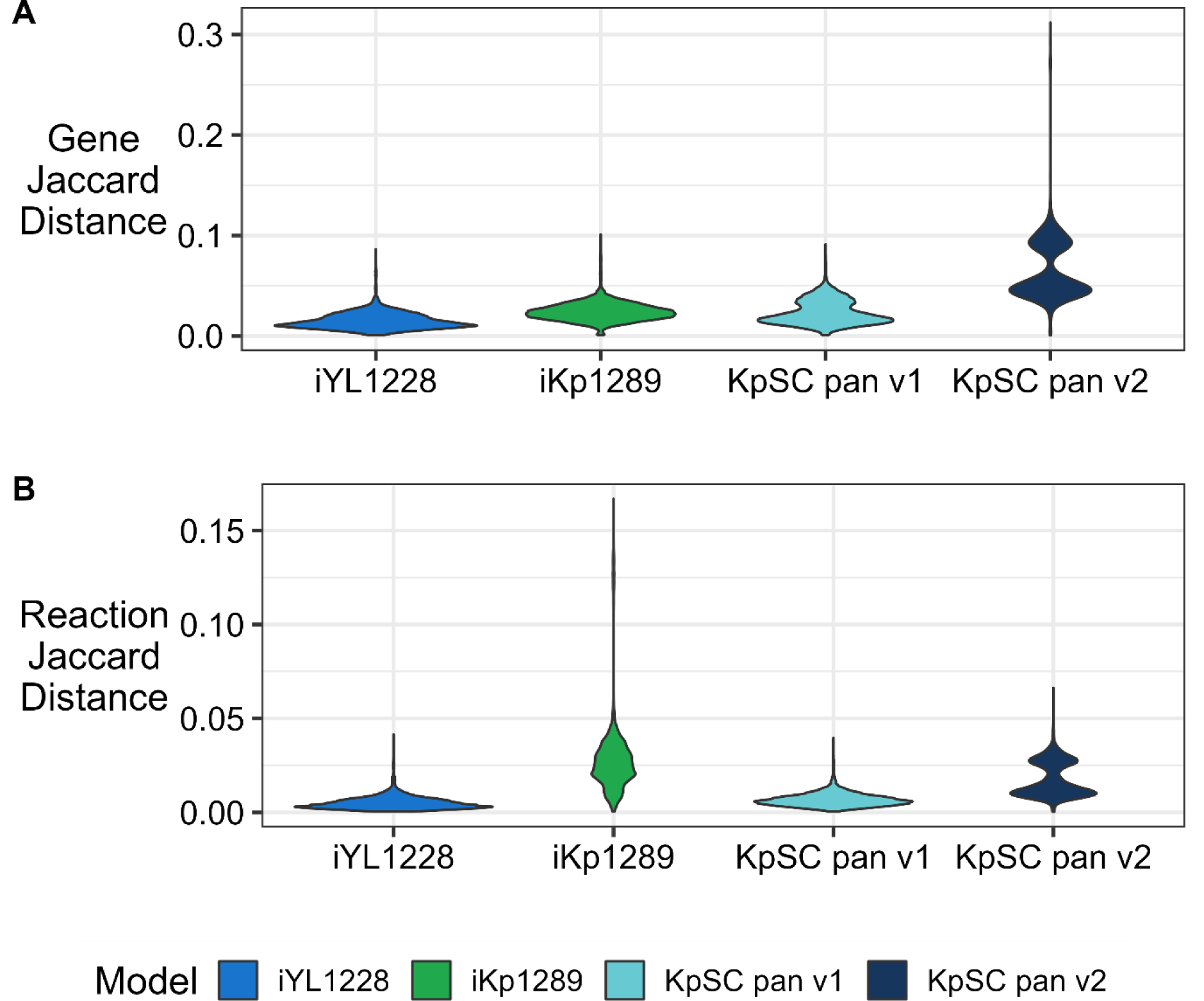
Pairwise gene and reaction Jaccard distances among strain-specific genome-scale metabolic models (GSMMs) derived from different reference models. Jaccard distances were calculated based on gene **(A)** and reaction **(B)** contents of pairs of strain-specific GSMMs derived from each of the iYL1223, iKp1289, KpSC pan v1 and KpSC pan v2 reference models. Kruskal-Wallis H-test indicated that the distributions were different (Genes: p < 2.2e-16, d.f. = 2,391 and Reactions: p < 2.2e-16, d.f. = 10,994). Pairwise comparisons indicated a significant difference between the distributions for KpSC pan v2 and all others (Mann-Whitney test; Genes & Reactions: p < 2.2e-16, Bonferroni correction threshold = 0.0167).

Distributions of pairwise reaction Jaccard distances (Figure 4B) supported the same trend as those identified for the gene distances (median pairwise distance 0.014 versus 0.004 (iYL1228), 0.006 (KpSC pan v1), p < 2.2×10^-16^ by Mann-Whitney test). However, the distribution for models derived from iKp1289 was unexpectedly broad (median = 0.025, range = 4.1×10^-4^ to 0.17). The longer skewed tail in the iKp1289 reaction distribution was driven by comparisons involving five genomes that were lacking two genes (VK055_1057 and VK055_0013) associated with 241 transport reactions. This disproportionately impacted the pairwise reaction Jaccard distances (Figure 4B). Within published GSMMs, these transport reactions are catalysed by the non-specific porins OmpK35 (48,49), OmpK36 (50,51), OmpK37 (52,53) and phosphoporin PhoE (54,55), that simulate the generic transport of hydrophilic molecules into the cell when the specific enzymes are unknown.

Transport of a molecule into the cell will not produce growth unless catalytic enzymes are present, thus sacrificing true molecular accuracy to increase accuracy of simulated phenotypic outcomes. While these porins are treated in the same way in all of the reference models, the gene sequences included in the iKp1289 model were not 100% conserved in our KpSC collection, inhibiting their detection and inclusion in the corresponding GSMMs when the default ortholog identity threshold was applied (80%).

As noted above, all KpSC are expected to grow in M9 minimal media supplemented with glucose, and we showed that all but four GSMMs generated using KpSC pan v2 as a reference were able to successfully simulate growth in these conditions. A single additional GSMM generated using the KpSC pan v1 reference was also unable to simulate growth (Table S1). In contrast, 71.0% (360/507) and 38.3% (194/507) GSMMs generated using the iYL1228 and iKp1289 reference models, respectively were unable to simulate growth in M9 plus glucose (Table S1). This finding reinforces the notion that single-strain models do not represent sufficient metabolic diversity for use as reference models in population analyses, either because they lack key metabolic pathways that are essential components of metabolism in a subset of the population and/or because they lack the allelic diversity that enables robust ortholog detection. In any case, in order to support further analysis of growth predictions, we used gap filling to identify and add missing reactions required to simulate growth in minimal conditions (13,14), restoring positive growth predictions for all models (excluding the known auxotrophs discussed above).

We compared the proportion of predicted core growth phenotypes (≥95% of isolates), accessory growth (<95% of isolates) and conserved no growth phenotypes for the collections of GSMMs generated using each of the reference models. Use of the KpSC pan v2 reference resulted in the highest proportion of accessory growth (9.2% versus 2.8%, 4.8% and 3.5%, Figure 3), with sulfur and phosphorus accessory growth phenotypes predicted only for models generated using the KpSC pan v2 reference. In addition, the proportions of conserved no growth phenotypes predicted by models generated with the KpSC pan v2 reference were lower than for those generated with its direct predecessor (KpSC pan v1, 44.6% versus 27.2%, respectively), demonstrating a higher number of completed metabolic pathways for simulatable growth sources.

### GSMMs derived from KpSC pan v2 are highly accurate

We compared the accuracy of GSMMs generated using each of the four reference models (KpSC pan v2 and (14,15,18)) for growth phenotype predictions in aerobic and anaerobic conditions, for a subset of 37 KpSC isolates with phenotypic growth data for 190 carbon substrates (5). For anaerobic conditions, we excluded *K. pneumoniae* MGH 78578 for which the data were considered spurious due to a high number of substrates that were not able to support growth, including those that are well characterised and known to generally support *Klebsiella* sp. growth in anaerobic conditions (e.g. glucose (56) and lactose (57)). The specific set of substrates that can be simulated is dependent on the reference model from which each GSMM was derived, and hence in order to facilitate fair comparisons we calculated accuracy metrics for multiple distinct sets of substrates: 89 substrates that were common to all GSMMs, 91 substrates common to GSMMs derived from iYL1228, KpSC pan v1 and KpSC pan v2, 105 substrates common to GSMMs derived from KpSC pan v1 and KpSC pan v2, and finally the full set of 124 substrates that could be simulated by GSMMs derived from KpSC pan v2 only (Figure 5 & Table 2). The phenotype results contain three substrates (L-lysine (58), glycine (59) and ethanolamine (60)) for which utilisation by KpSC is known or expected to be tightly regulated, meaning the standard growth conditions are not expected to stimulate growth. For example, L-lysine utilisation is expected to be universal in the KpSC (61,62) and is correctly predicted as such for GSMMs derived from KpSC pan v2 (Table S6), but the required decarboxylase is only active in acidic conditions (58) resulting in no growth in the phenotype data (5). As GSMMs do not consider such gene regulation or slow growth rates (11), we have excluded L-lysine, glycine and ethanolamine from the accuracy metrics presented below.

**Figure 5:**
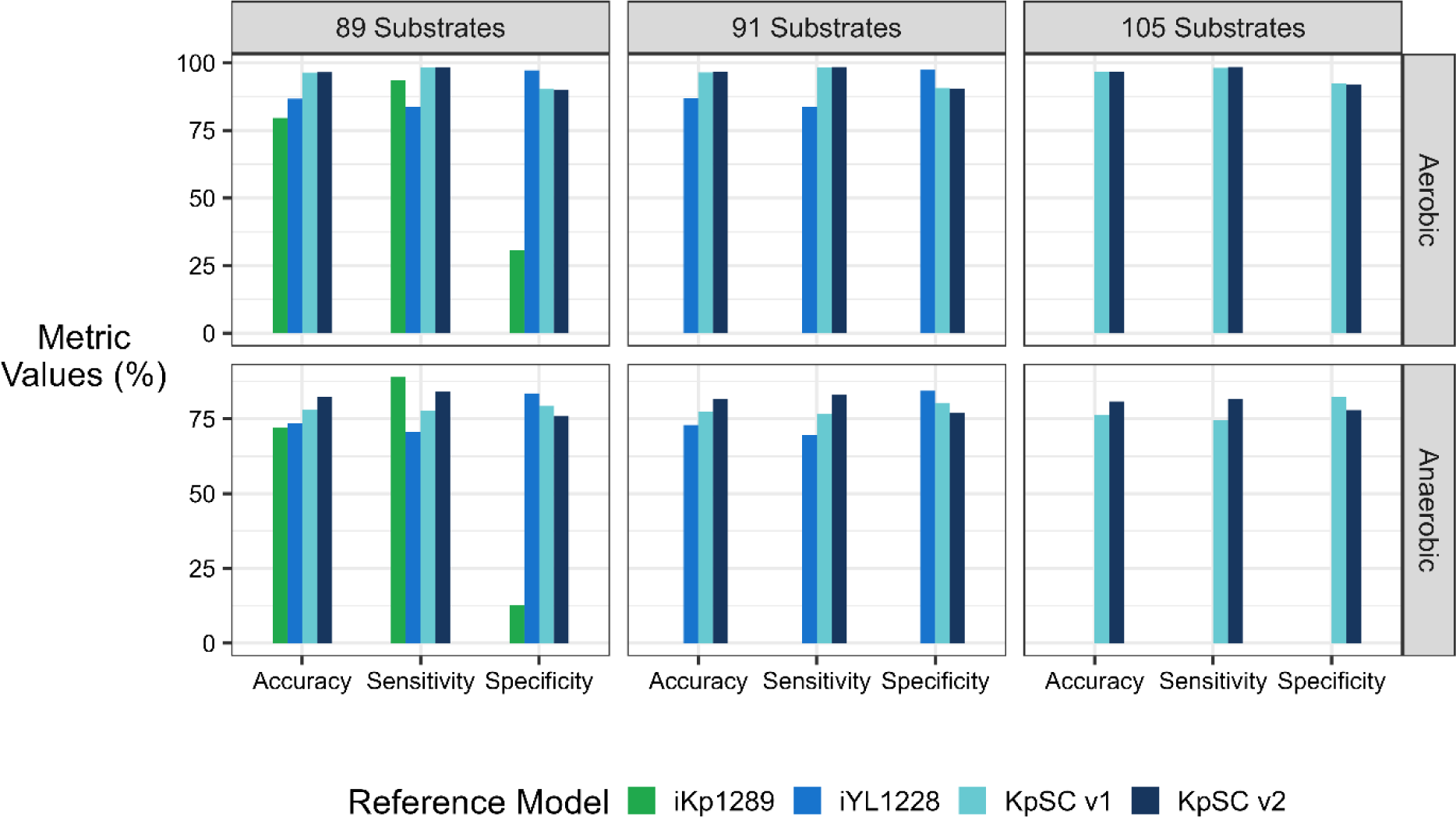
Comparable growth prediction accuracies for strain-specific genome scale metabolic models derived from iKp1289, iYL1228, KpSC pan v1 and KpSC pan v2 references. Accuracies, sensitivities and specificities represent combined values for carbon substrate growth predictions compared to true phenotypes generated on the Biolog platform (plates PM1 and PM2) for 37 KpSC isolates. Aerobic conditions are presented in the first row (raw values in Table S6) and the anaerobic conditions in the second row (raw values in Table S7). Left panels comprise data for the set of 89 substrates supported by all four models. Middle panels comprise data for the same 89 substrates plus two additional substrates supported by iYL1228, KpSC pan v1 and KpSC pan v2 (91 substrates in total). Right panels comprise data for the same set of 91 substrates plus an additional 14 substrates supported by only KpSC pan v1 and KpSC pan v2. Table 2 shows metrics calculated for the total number of substrates supported by each reference model.

**Table 2:** Overall model accuracy for aerobic and anaerobic carbon sources using the four *Klebsiella* reference models.

|  |  | iKp1289 | iYL1228 | KpSC pan v1 | KpSC pan v2 |
| --- | --- | --- | --- | --- | --- |
|  | Number of Substrates Tested<sup>1</sup> | 95 | 91 | 105 | 124 |
| Aerobic | Overall Accuracy (%) | 77.92 | 86.87 | 96.65 | 95.38 |
|  | Strain-Specific Range (%) | 72.63 - 82.11 | 81.32 - 91.21 | 92.38 - 98.10 | 87.90 - 97.58 |
|  | True Positives | 2480 | 2174 | 2817 | 3245 |
|  | True Negatives | 259 | 751 | 938 | 1131 |
|  | False Positives | 525 | 20 | 76 | 155 |
|  | False Negatives | 251 | 422 | 54 | 57 |
|  | Sensitivity (%) | 90.81 | 83.74 | 98.12 | 98.27 |
|  | Specificity (%) | 33.04 | 97.41 | 92.50 | 87.95 |
| Anaerobic | Overall Accuracy (%) | 70.23 | 72.92 | 76.38 | 78.76 |
|  | Strain-Specific Range (%) | 43.16 - 81.05 | 59.34 - 84.62 | 61.90 - 86.67 | 59.68 - 87.90 |
|  | True Positives | 2272 | 1758 | 2129 | 2594 |
|  | True Negatives | 130 | 631 | 758 | 922 |
|  | False Positives | 664 | 117 | 162 | 227 |
|  | False Negatives | 354 | 770 | 731 | 721 |
|  | Sensitivity (%) | 86.52 | 69.54 | 74.44 | 78.25 |
|  | Specificity (%) | 16.37 | 84.36 | 82.39 | 80.24 |

<sup>1</sup>Represents the total number of substrates supported by the model for which matched aerobic and anaerobic phenotype data generated on the Biolog platform were available. Note that substrates supported by iYL1228 are a subset of those supported by KpSC pan v1, which are a subset of those supported by KpSC pan v2. Substrates supported by iKp1289 are an independent but overlapping set.

GSMMs derived from the KpSC pan v2 reference showed the highest overall accuracy for each set of comparable growth predictions (96.6% aerobic & 82.3% anaerobic for n=89 substrates, 96.6% aerobic & 81.6% anaerobic for n=91 substrates, 96.8% aerobic & 80.8% anaerobic for n=105 substrates, Figure 5). This is followed by GSMMs derived from KpSC pan v1 (96.5% aerobic & 78.1% anaerobic for n=89 substrates, 96.5% aerobic & 77.4% anaerobic for n=91 substrates, 96.7% aerobic & 76.4% anaerobic for n=105 substrates, Figure 5) and those derived from iYL1228 (86.7% aerobic & 73.4% anaerobic for n=89 substrates, 86.9% aerobic & 72.9% anaerobic for n=91 substrates, Figure 5). GSMMs derived from the iKp1289 reference resulted in the lowest overall accuracy (79.6% aerobic & 72.0% anaerobic for n=89 substrates, Figure 5), and were associated with particularly low sensitivity (30.7% aerobic & 12.7% anaerobic, Figure 5). In contrast, there was a trend towards increasing sensitivity and decreasing specificity from GSMMs derived from iYL1228, to those derived from KpSC pan v1, and those derived from KpSC pan v2, implying that as more metabolic content is added to successive versions of these reference models, the ability to accurately predict growth increases at the cost of lower accuracy in predicting no-growth (Figure 5). This behaviour is to be expected, because false positive predictions are assumed to result from regulation that may be preventing growth *in vitro* (not considered by GSMMs) whereas false negative predictions imply that genetic and/or metabolic information is missing from the model (11).

The final accuracies calculated for all 124 substrates that could be simulated by KpSC pan v2 derived GSMMs was 95.4% and 78.8% in aerobic and anaerobic conditions respectively (Table 2), representing a decrease of 1.2% and 2.0% compared to those calculated for the n=105 substrate subset (Tables S6 & S7). Consequently, the final accuracy estimate for KpSC pan v2 derived GSMMs (124 substrates) is lower than those derived from KpSC pan v1 (105 substrates, Table 2), despite the superior accuracy of KpSC pan v2 in the comparative analysis (Figure 5). This has occurred because most of the substrates newly supported by KpSC pan v2 are less likely to support growth of KpSC strains and are therefore understudied. One such example is 2-Deoxy-D-Ribose for which the aerobic growth prediction accuracy was 27.0% (Table S6). In *Salmonella* (63), 2-Deoxy-D-Ribose is degraded by the 2-Deoxy-D-Ribose utilisation operon (*deoQKPX*), with the model encoding *deoK* (deoxyribokinase) and *deoP* (inferred deoxyribose permase). However, neither *deoK* or *deoP* have been fully characterised and cannot be accurately distinguished from ribokinases (*rbsK*) or L-fucose symporters (*fucP*) respectively at the protein level (37) (both are 100% conserved in our KpSC isolate collection). As the correct genes could not be identified, the associated GPR reactions were left empty (i.e. all GSMMs derived from the reference can inherit the reaction), resulting in an incorrect core growth prediction. In spite of this, we have elected to retain the pathway in the model because there is sufficient evidence that *Klebsiella* can utilise this substrate and the pathway is known (63). The model can easily be updated when further information is available to identify the true sequences.

As previous phenotype experiments do not contain all possible substrates that can be newly simulated by the KpSC GSMMs, we performed independent growth assays for six substrates, for which the estimated accuracies ranged from 32.4% to 90.5% (Table 3).

**Table 3:** Summary of additional phenotypic validation experiments for KpSC pan v2.

| Substrate<sup>1</sup> | Predicted Growth<sup>2</sup> | TP | TN | FP | FN | Accuracy |
| --- | --- | --- | --- | --- | --- | --- |
| 2-Aminoethylphosphonate | 31/37 | 6 | 6 | 25 | 0 | 32.4% |
| Beta-Alanine | 37/37 | 17 | 0 | 20 | 0 | 46.0% |
| Allantoin | 3/37 | 2 | 31 | 1 | 3 | 89.2% |
| L-ectoine | 5/41 | 1 | 34 | 4 | 2 | 85.4% |
| Phenylpropanoate | 10/42 | 7 | 31 | 2 | 2 | 90.5% |
| Uracil | 37/37 | 28 | 0 | 9 | 0 | 75.7% |

<sup>1</sup>Substrates were not present on Biolog PM1 / PM2 plates, and were tested via an independent culture approach in aerobic conditions only (See Methods).
<sup>2</sup>Isolates tested include 37 KpSC for which Biolog microarray data were available. Two phenotypes were predicted as rare among this set of 37 KpSC, and hence additional isolates for which growth was predicted were selected from our collection; n=4 available for L-ectoine, n=5 for Phenylpropanoate (see Table S5).

Phenylpropanoate utilisation was the most accurately predicted, with high sensitivity (77.8%) and specificity (93.9%, Table 3). Growth on L-ectoine, allantoin and uracil were also predicted with accuracies >75%; however, we note that the total number of isolates predicted to be able to utilise L-ectoine and/or allantoin was low (≤5). This is to be expected, with allantoinase (required for allantoin utilisation) found to be present in only 5% of *K. pneumoniae* and 50% of *K. quasipneumoniae* (8). The low accuracies for 2-Aminoethylphosphonate (32.4%) and beta-alanine (46.0%) may be explained by mechanisms not taken into account by GSMMs (11), such as gene regulation or experimental conditions. In the case of 2-Aminoethylphosphonate usage, metabolism in other *Enterobacteriaceae* species only occurs in conditions of phosphate starvation (64), while beta-alanine metabolism may require nitrogen starvation (65). Alternatively, the discrepancies may be due to limitations with using cellular growth (OD600) instead of tetrazolium dye as a proxy of cellular respiration to determine substrate utilisation. In any case, when these six additional substrates were included, the overall accuracy for aerobic growth conditions for KpSC pan v2 derived GSMMs was 94.3% (Sensitivity = 98.1% & Specificity = 85.4%).

## 5. Discussion

We present a highly curated and accurate KpSC pan metabolic model, capturing the highest number of distinct metabolic reactions of any pan-model published to-date for any bacterial taxa (66–69). We show that KpSC pan v2 successfully captures a greater proportion of variable metabolic traits than its predecessors (Figures 3 & 4) and can be used as a reference to generate strain-specific GSMMs with superior completeness (in terms of metabolic pathways supporting core glucose metabolism) and accuracy (for growth phenotype predictions, Figure 5). When compared to growth phenotype data, strain-specific GSMMs derived from KpSC pan v2 were associated with 95.4% and 78.8% accuracy for aerobic and anaerobic conditions, respectively (n=124 substrates, Table 2). However, six additional substrates tested via our in-house OD600 approach in aerobic conditions only, were associated with a wider range of prediction accuracies (32.4% to 90.5%, Table 3). The lower accuracy predictions arose from higher number of false positives (sensitivity = 33.3% to 100%, specificity = 0% to 96.9%, Table 3), likely from a combination of methodological limitations and regulatory effects that are not accounted for by GSMMs, the latter being a key caveat of *in silico* growth predictions that has been recognised previously (11).

The KpSC pan v2 model represents 507 individual KpSC isolates that were selected based on the availability of existing curated GSMMs (6,14,15,18), and to represent broad population diversity. The previously published GSMMs included 37 isolates with matched phenotype data (5) that represented all KpSC subspecies. To further diversify this set we included 452 isolates from published clinical diversity collections (16,70,71), which have been extensively characterised and for which the physical isolates were available in our laboratory. These collections represent clinical KpSC isolated from an Australian hospital network over a 12 month period (16), plus isolates from rectal swabs of patients in select wards (70,71). Isolates were included regardless of their antibiotic resistance status (23.7% multidrug resistant), and were genetically diverse (16,70,71). Overall, the 507 isolates represented in KpSC pan v2 comprise 268 unique STs and include representatives of 12 of 13 globally disseminated problematic clones described in the literature (7) (lacking clonal group 307, Table S1). However, we acknowledge that this collection may not fully represent the true global KpSC population, which comprises hundreds to thousands of unique clones (16,43,71). Additionally, most isolates were either human clinical or gut carriage isolates (97.5% of isolates), with only a small proportion from environmental sources or other host-associated environments (Table S1). Hence, it is likely that the model excludes metabolic traits specific to non-human niches. Furthermore, the inclusion of metabolic traits is limited by the bounds of the existing biochemical and biological knowledge bases, which are undoubtedly biassed towards common metabolic processes. As a consequence, we anticipate that rare metabolic traits are underrepresented in the model, and accordingly the relative proportion of accessory versus core reactions (8.8% accessory & 91.2% core) captured in the model is far smaller than that estimated for the total KpSC gene pool (8,16).

The total pangenome derived from the 507 isolates included in this work comprised 34,664 unique genes and the final curated KpSC pan v2 model includes just 2,403 of these (∼7%). However, a previous automated analysis indicated that up to 37% of the KpSC pangenome encodes proteins involved with metabolic processes (8), further highlighting the limitations in the underlying knowledge bases that inform metabolic models (in particular where reaction stoichiometries are not defined). In addition, it is important to consider the types of evidence available to support the inclusion of genes and reactions in the model, which in most cases is limited to sequence homology and/or simple growth phenotype data (evidence score 2, 81.9%). Novel proteomic and/or protein structural data generated specifically for KpSC isolates may enable the inclusion of additional reactions with greater confidence, as demonstrated for construction of the *Salmonella enterica* pan model (72). Targeted gene-knockout and/or gene expression analyses conducted in aerobic and anaerobic conditions could also shed new light on the associated metabolic differences and inform improvements for anaerobic growth predictions.

Despite its limitations, KpSC pan v2 represents a significant resource for the *Klebsiella* research community, both as a source of curated KpSC-specific metabolic information and as a reference model from which highly accurate strain-specific GSMMs can be readily derived. The model will be updated as additional biochemical data and evidence become available. It is freely available for use, reuse and adaptation under open access licence and we welcome contributions from the community. We anticipate that KpSC pan v2 will inform studies of KpSC virulence and antimicrobial resistance, as well as KpSC ecology (e.g. within the human gut microbiota), supporting the development of novel therapeutics and containment strategies targeting KpSC. Furthermore, we hope that our data highlighting the benefits of the pan-model approach will inspire similar works for other prominent bacterial pathogens, particularly those known to harbour high levels of genomic diversity such as *E. coli*, *Pseudomonas aeruginosa* and *Acinetobacter baumannii*.

## 6. Author statements

### 6.1 Author contributions

1. Conceptualisation: KLW, KEH & JMM
2. Data Curation: HBC
3. Formal Analysis: HBC
4. Funding Acquisition: HBC, KLW, KEH, SB, JMM
5. Investigation: HBC, SB, VP, SLF
6. Methodology: HBC & KLW
7. Project Administration: KLW
8. Resources: KLW, KEH & SB
9. Software: HBC & BV
10. Supervision: KLW, JH & KEH
11. Validation: HBC, BV, VP & SLF
12. Visualisation: HBC
13. Writing - Original Draft: HBC & KLW
14. Writing - Review & Editing: All Authors

## Acknowledgements

The authors would like to thank Kalani Paranagama (Monash University) for their discussions and advice regarding this project.

