## Supplementary Figures for "A validated pangenome-scale metabolic model for the *Klebsiella pneumoniae* species complex"

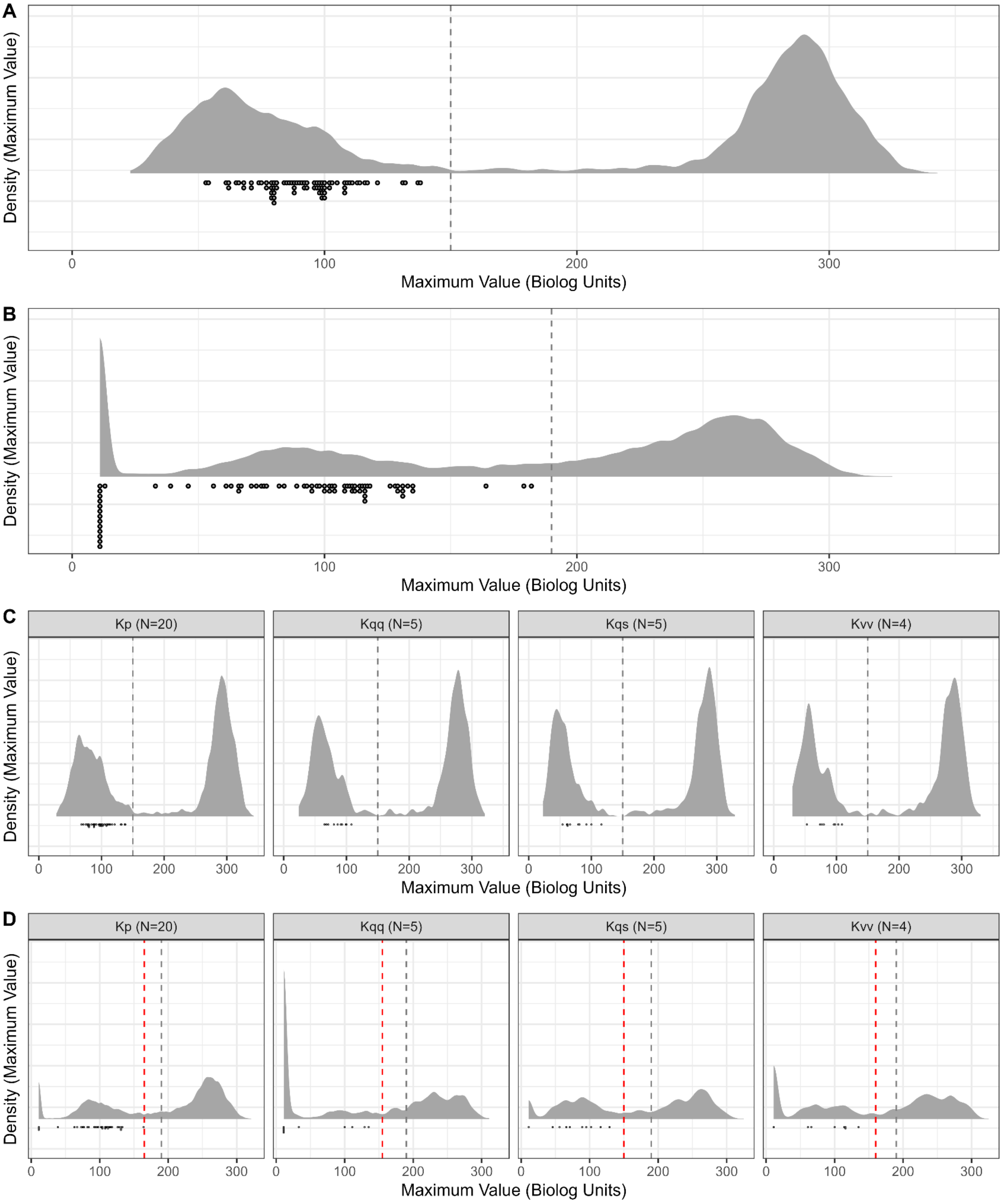
A
Density (Maximum Value)
0
100
200
300
Maximum Value (Biolog Units)
B
Density (Maximum Value)
0
100
200
300
Maximum Value (Biolog Units)
C
Kp (N=20)
Kqq (N=5)
Kqs (N=5)
Kvv (N=4)
Density (Maximum Value)
0
100
200
300
Maximum Value (Biolog Units)
D
Kp (N=20)
Kqq (N=5)
Kqs (N=5)
Kvv (N=4)
Density (Maximum Value)
0
100
200
300
Maximum Value (Biolog Units)

**Figure S1: Maximum growth value distributions and empirical thresholds for 37 KpSC isolates subjected to phenotyping on the Biolog microarray.** Distribution of maximum values (Biolog units) for each of 190 substrates (plates PM1 and PM2) tested for each of 37 KpSC isolates in **(A)** aerobic and **(B)** anaerobic conditions. Species specific distributions of maximum values for the same set of 190 substrates tested for 5 - 20 isolates each (as indicated) in aerobic **(C)** and anaerobic **(D)** conditions. Species represented by only a single isolate are not shown. Black dots beneath the distributions are the negative control values (no isolate/nutrient controls), which should always be classified as no growth. Black dotted lines represent the empirical species complex growth thresholds for aerobic (5) and anaerobic conditions. Red lines represent the species-specific thresholds for anaerobic data; *K. pneumoniae* (Kp, x=165), *Klebsiella quasipneumonise* subsp *quasipneumoniae* (Kqq, x=155), *Klebsiella quasipneumonise* subsp *similipneumoniae* (Kqs, x=150) and *Klebsiella variicola* subsp *variicola* (Kvv, x =160).

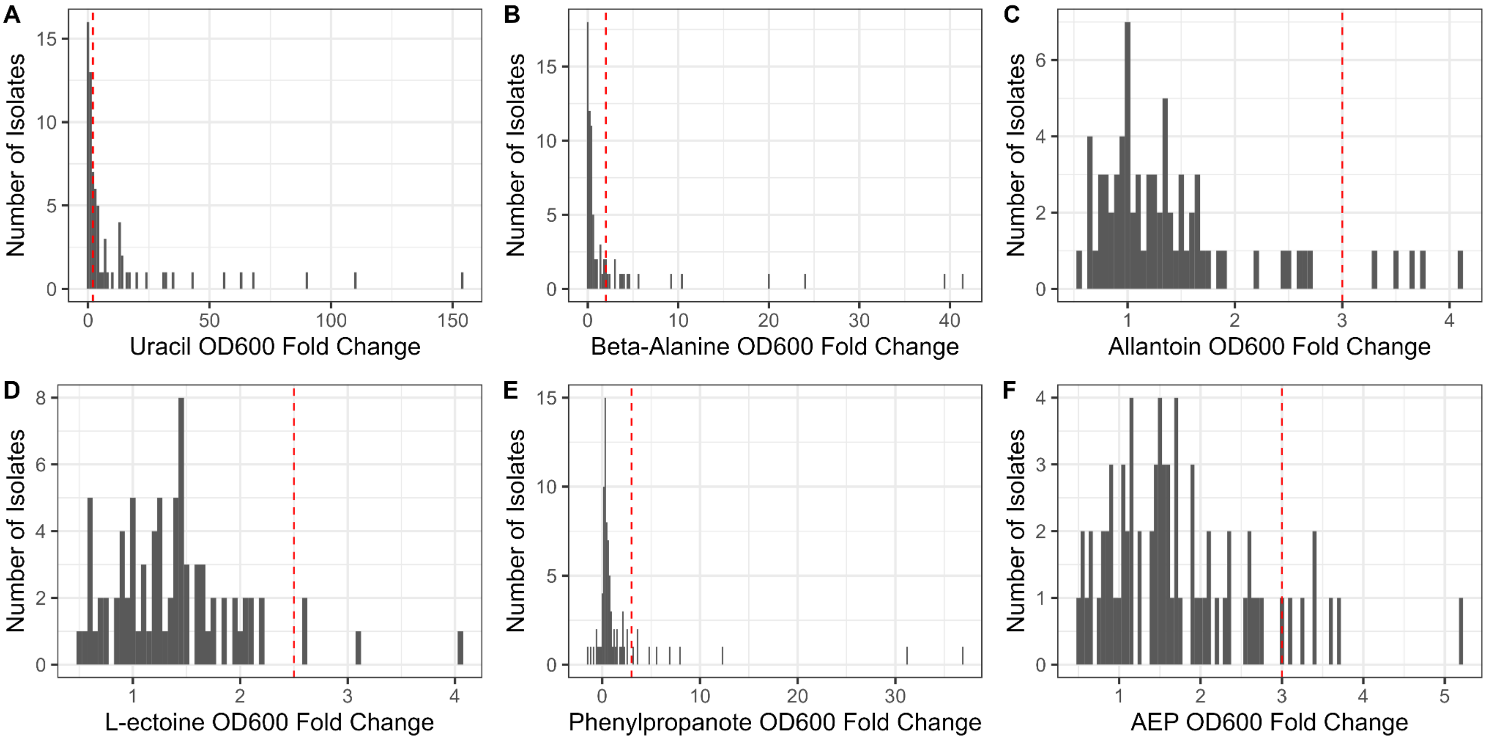


**Figure S2: Growth distributions and empirical thresholds for isolates cultured in minimal media plus a single carbon source.** Bars show mean $OD_{600}$ fold-change compared to the substrate-negative control for the same isolate (n=3 technical replicates, see Methods). Red dotted lines indicate growth thresholds defined from the empirical distributions: **(A)** uracil, growth ≥ 2.0; **(B)** beta-alanine growth ≥ 2.0; **(C)** allantoin, growth ≥ 3.0; **(D)** L-ectoine, growth ≥ 2.5; **(E)** phenylpropanoate, growth ≥ 3.0; **(F)** 2-Aminoethylphosphonate (AEP), growth ≥ 3.0.

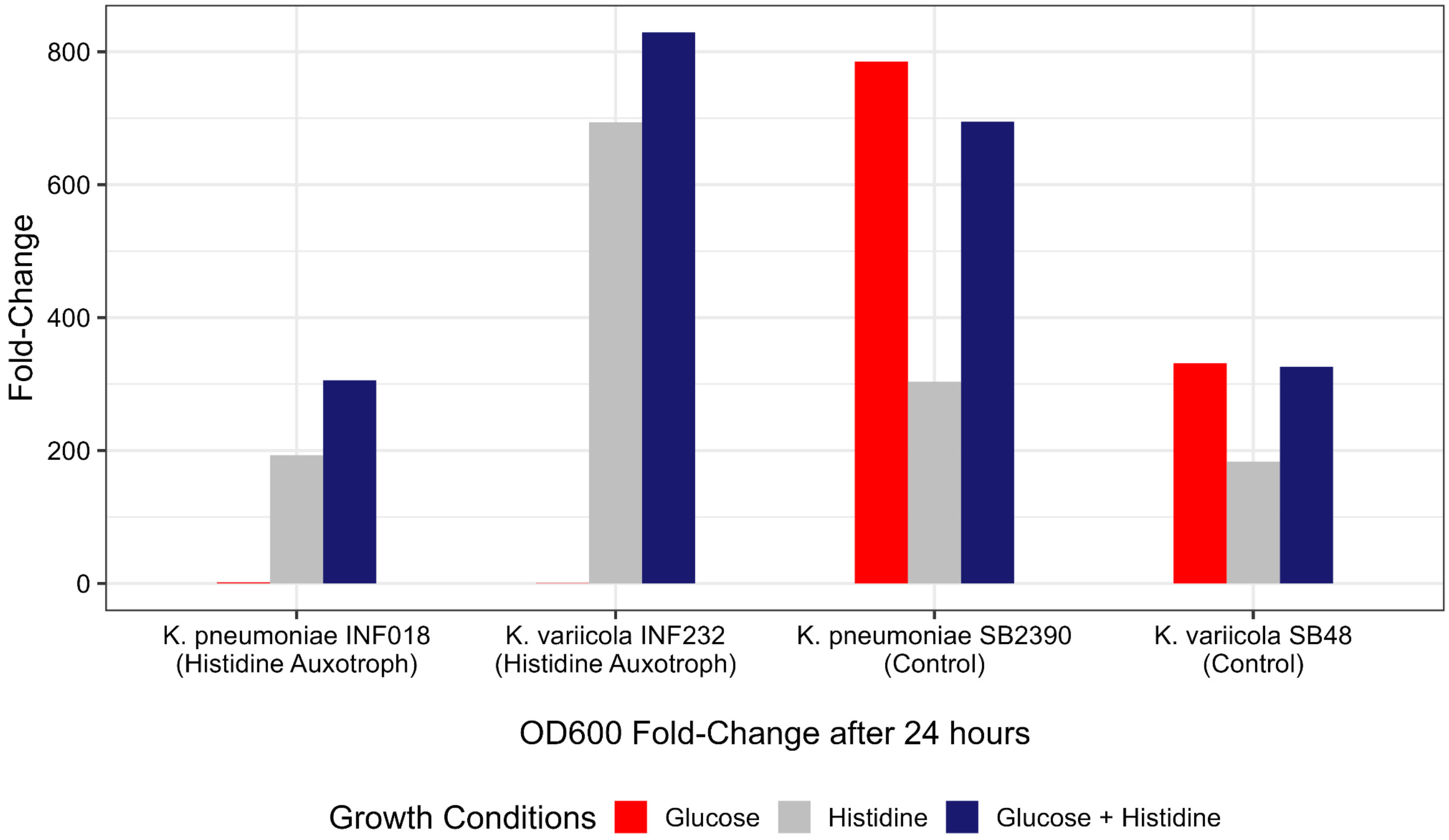


**Figure S3: Confirmation of histidine auxotrophy.** Suspected auxotrophs (*K. pneumoniae* INF018 and *K. variicola* INF232) and control isolates (*K. pneumoniae* SB2390 and *K. variicola* SB48) were cultured in minimal media with/without glucose and/or histidine (as indicated in legend). Bars show mean $OD_{600}$ fold-change at 24 hours compared to the substrate-negative control for the same isolate (n=3 technical replicates, see Methods). Control isolates showed clear evidence of growth in all conditions (fold change ≥ 2), albeit at a lower rate in the absence of glucose (grey bars) which we attribute to comparatively inefficient metabolism of histidine as a carbon source. In contrast, the suspected histidine auxotrophs were able to grow only when exogenous histidine was available in the growth media.

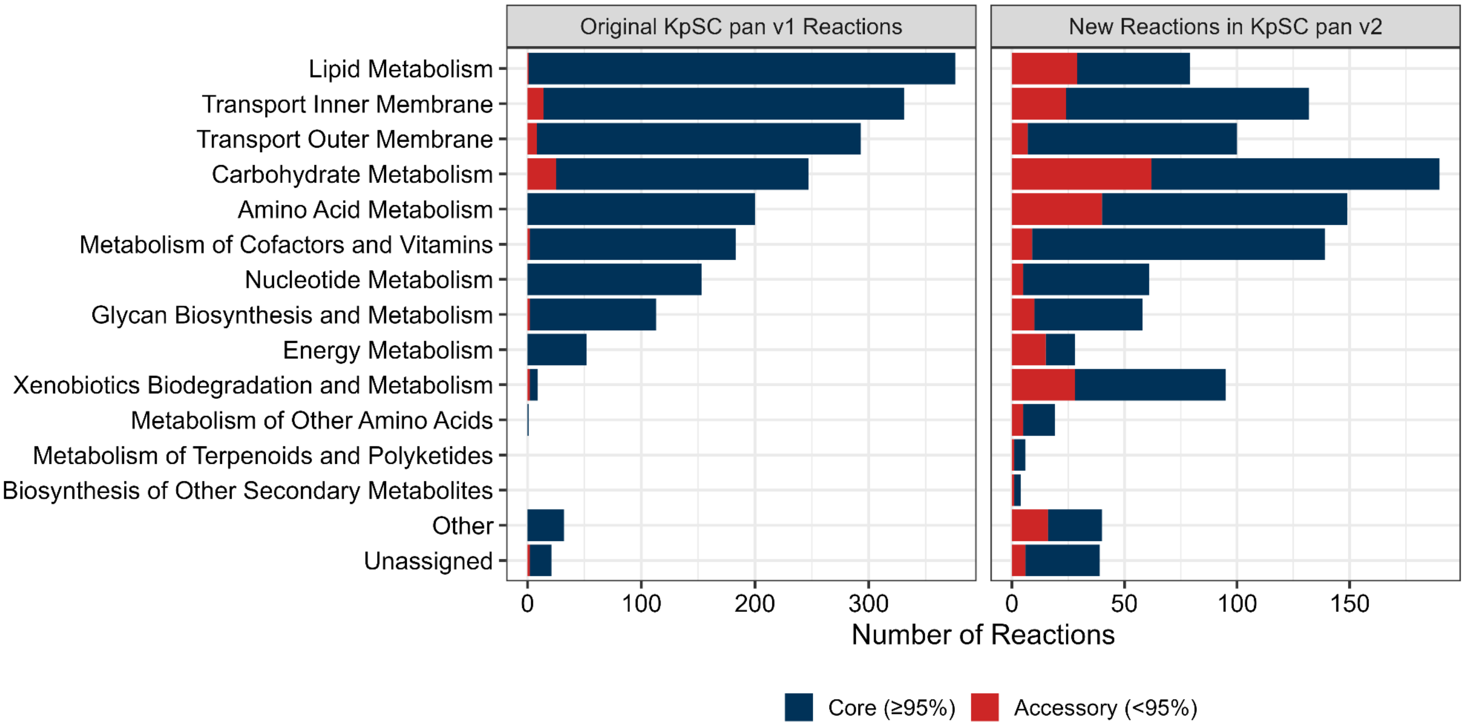


**Figure S4: Population conservation of reactions in KpSC pan v1 and KpSC pan v2.** The left panel refers to the reactions that were originally present in KpSC pan v1 (15), whereas the right panel refers to reactions that were newly added to KpSC pan v2. Core (dark blue) and accessory (red) reaction subsystems are defined as reactions present in ≥95% and <95% of 505 strain-specific genome scale metabolic models, respectively. Other refers to reaction subsystems that were not present on KEGG whereas unassigned refers to reactions that had no subsystem classification assigned. Transport reactions are not part of the KEGG subsystems, so these were grouped by whether they moved metabolites across the inner or outer membrane.
